# Using Neural Style Transfer to study the evolution of animal signal design: A case study in an ornamented fish

**DOI:** 10.1101/2023.03.13.532060

**Authors:** Yseult Héjja-Brichard, Kara Million, Julien P Renoult, Tamra C Mendelson

## Abstract

The sensory drive hypothesis of animal signal evolution describes how animal communication signals and preferences evolve as adaptations to local environments. While classical approaches to testing this hypothesis often focus on preference for one aspect of a signal, deep learning techniques like generative models can create and manipulate stimuli without targeting a specific feature. Here, we used an artificial intelligence technique called neural style transfer to experimentally test preferences for color patterns in a fish. Findings in empirical aesthetics show that humans tend to prefer images with the visual statistics of the environment because the visual system is adapted to process them efficiently, making those images easier to process. Whether this is the case in other species remains to be tested. We therefore manipulated how similar or dissimilar male body patterns were to their habitats using the Neural Style Transfer (NST) algorithm. We predicted that males whose body patterns are more similar to their native habitats will be preferred by conspecifics. Our findings suggest that both males and females are sensitive to habitat congruence in their preferences, but to different extents, requiring additional investigation. Nonetheless, this study demonstrates the potential of artificial intelligence for testing hypotheses about animal communication signals.

**Highlights:**

– Neural style transfer was used to test preferences for patterns in a colorful fish
– Results trend in support of the processing bias hypothesis, a component of sensory drive
– This study demonstrates the use of generative AI for animal behavioral studies

## Introduction

Animals are characterised by a vast array of colours and patterns that evolve due to both natural and sexual selection: animals need to be able to hide from predators and to be visible to potential mates. In both contexts, the habitat plays a crucial role in the evolution of colour patterns. In the case of predation, avoidance may involve background-matching (Merilaita & Lind, 2005) or disruptive colouration (Cuthill et al., 2005; Stevens et al., 2008), two camouflage strategies that rely on matching or contrasting characteristics (colours, patterns) of the visual environment, respectively (see e.g., Cuthill et al., 2016). In the case of mating, for an animal to successfully convey a courtship message, it must be detectable by the receiver, meaning an individual must stand out from its background. However, based on theoretical and empirical results in human visual preferences (Menzel et al., 2015), and correlative studies in animals (Hulse et al., 2020), authors have proposed that sexual signals could also be more attractive if their patterning matches that of the background (Renoult & Mendelson, 2019). This hypothesis remains to be tested empirically in animals.

For attracting a mate, conspicuous signalling can be achieved with a broad diversity of colours and patterns, and understanding the drivers of this diversity in sexual signals has been a central focus of evolutionary biology. One of the most influential hypotheses is sensory drive (Endler, 1992; Endler & Basolo, 1998; Seehausen et al., 2008). Based largely on signal detection theory, sensory drive suggests that the need to be optimally conspicuous leads to the evolution of different signals in different environments. Most evidence supporting this hypothesis comes from studies that investigate the link between one feature of an animal signal design, such as colouration, and the corresponding physical characteristic in the environment, like the colour of ambient light. The main prediction of sensory drive is that the detectability of a signal should be favoured to have a higher signal-to-noise ratio. A compelling example is the case of the colourful dewlap of Anolis lizards, which is used for courtship and competition signalling. Anolis species in bright habitats tend to have red or orange dewlaps, while species in darker-lit habitats tend to have white or yellow dewlaps, both maximizing visibility in their respective habitats (Fitch & Hillis, 1984; Fleishman, 1992; Fleishman et al., 2022).

However, to attract a mate, a signal must be more than detected. A robust finding in empirical aesthetics is that the subjective ease with which a signal is processed by the brain is a strong determinant of how attractive that signal will be; this is the fluency theory of aesthetics, which predicts that a stimulus that is easy to process (depending on both the stimulus’ intrinsic properties and the observer’s perceptual processing of that stimulus) will trigger positive affect for its observer (Fantz, 1957; Munar et al., 2015; Reber et al., 2004; Renoult et al., 2016). In addition, neuroscientists have extensively documented that perceptual systems are adapted to efficiently process natural scenes and their spatial statistics (Olshausen & Field, 1997; Simoncelli & Olshausen, 2001). As a consequence, signal designs that imitate the spatial statistics of natural scenes should be processed more efficiently, because they match the sensitivity, or tuning, of neurons that are adapted to those habitats. Combined with the fluency theory of aesthetics, this prediction suggests that signal designs that imitate the spatial statistics of natural scenes should also generate a pleasant feeling of fluency (Geller et al., 2022) and thus they are likely attractive. Combining the fluency theory of aesthetics with sensory drive, Renoult & Mendelson (2019) proposed the concept of a processing bias and suggested that this mechanism could explain the attractiveness and evolution of complex signal designs in both human and animal communication.

One way to test the attractiveness of sexual signals that mimic the patterns of natural habitats is to use the Fourier slope, which corresponds to the distribution of spatial frequencies (i.e. luminance distribution) in pattern images. In camouflage studies, the Fourier slope has been shown to reliably predict the degree of background matching in cuttlefish and octopus (Zylinski et al., 2011; Josef et al., 2012). Similarly, the Fourier slope can be used to determine whether background matching plays a role in attractiveness. For instance, jumping spiders prefer abstract patterns that more closely match the Fourier sope of terrestrial habitats (Hardenbicker & Tedore, 2023). However, the Fourier slope only captures one aspect of a visual pattern: the scale invariance in the distribution of luminance contrasts. Other features could be equally or even more important in explaining how background matching can increase visual attractiveness.

Testing preferences for complex designs and for designs that do not yet exist requires new approaches. Tools from artificial intelligence and in particular from deep learning represent a valuable complementary approach. Generative models, such as variational autoencoders (VAEs; Kingma & Welling, 2019) and generative adversarial networks (GANs, Goodfellow et al., 2014; 2020) can be used to create or manipulate stimuli and study animal preferences without the need to target a specific feature. Exploiting the architecture of GANs, (Talas et al., 2020) developed CamoGAN to study the evolutionary arms race between the camouflage of a synthetic prey and its predator. Another example is CycleGAN (Zhu et al., 2020). This GAN-based algorithm can be trained to transform one image into a different category and could be used to model an intermediate between two phenotypes or between two species to study conspecific preferences. A complementary method, StyleGAN3 (Karras et al., 2021), can be used to modify specific features of an image. Another type of generative model is the (latent) diffusion model (Sohl-Dickstein et al., 2015; Ho et al., 2020; Song & Ermon, 2020). Such a model can take a sentence as input and turn it into an image (see e.g. (Ramesh et al., 2021; Nichol et al., 2022) and was made famous with the release of the DALL-E application by OpenAI. Diffusion models can thus be used to synthesize rich and controlled images to test specific hypotheses about signal preferences. Although they have interesting properties and can generate high-quality images, such models require a large number of images and extensive computer power to be trained, and they have a somewhat complex architecture. The use of APIs such as DALL-E can be employed as an alternative but offers little control to the user.

One tool of artificial intelligence that allows a high level of control over stimulus generation is the Neural Style Transfer (NST) algorithm developed by Gatys and colleagues (2015, 2017). The algorithm requires two input images, one for content and one for style, and aims to render the content image in the style of the style image. Using the architecture of a convolutional neural network (CNN), the NST algorithm extracts the content, namely the global arrangement of an image, such as the objects that constitute it, and the style, computing the Gram matrix of an image which corresponds to the texture information and local variations of luminance, of two images, respectively. The algorithm creates a third image that combines the content of one image with the style of the other (Figure 1). Importantly, the user can determine how much weight to put on the style or on the content to modulate the output image as desired. Moreover, the algorithm only requires two input images and no extra training on the user end, and several accurate implementations are ready to use along with tutorials.

**Figure 1.**
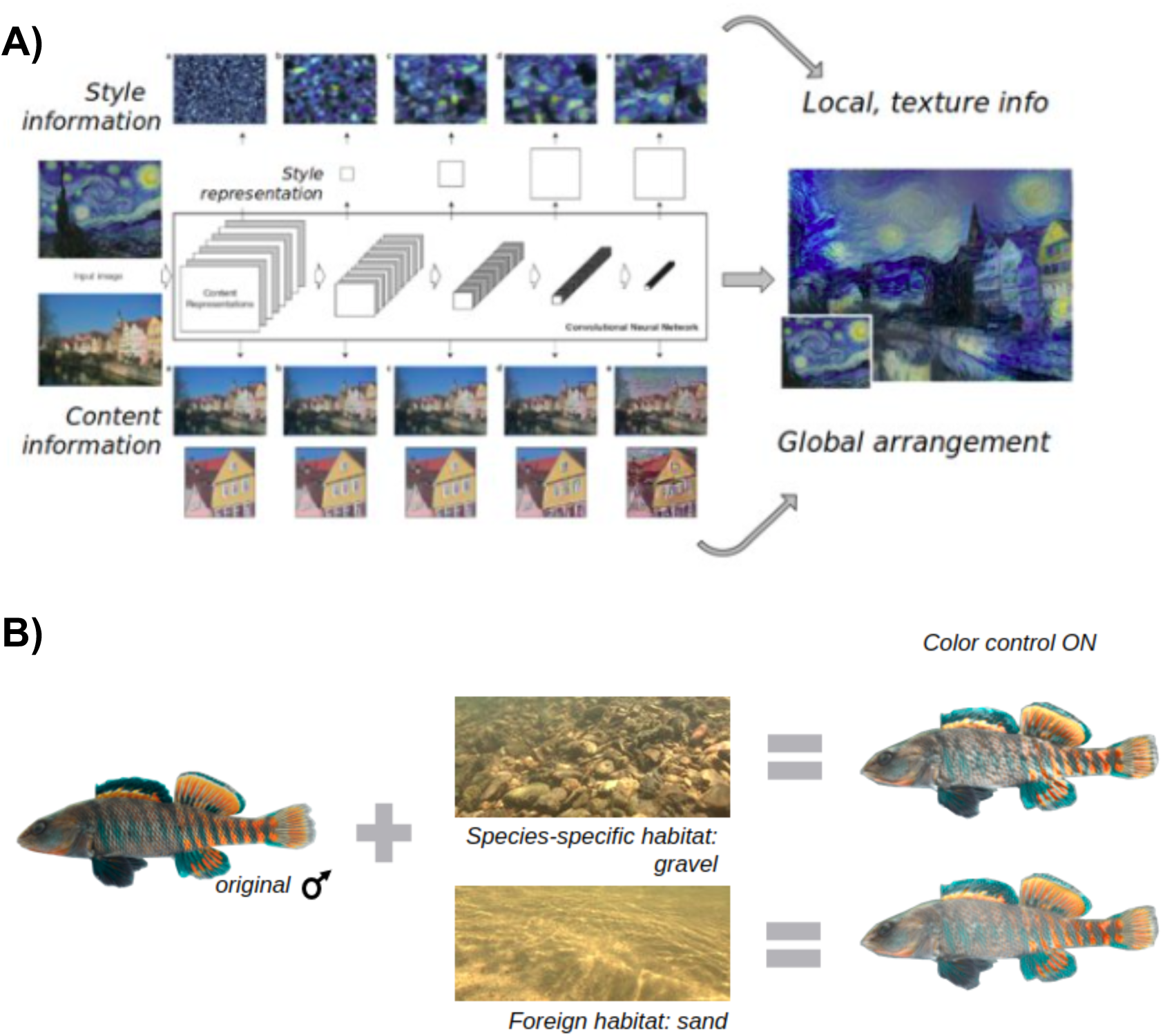
Neural Style Transfer. Panel A: The algorithm produces a new image by minimising the distance with both content and style images. The style information corresponds to local statistics of the image (texture, colour, paint strokes), while the content information corresponds to the global arrangement of an image (what the image is made of, e.g. the row of houses along the river). Adapted from Gatys et al., 2016. Panel B: Creation of the stimuli. A picture of an original fish (male individual) and a picture of the habitat (either gravel or sand) are fed to the Neural Style Transfer algorithm as content and style images, respectively. The algorithm extracts the style information via a Gram matrix while the content information corresponds to the usual feature layers of a CNN. The original colours of the fish were put back on the final image to make it look more naturalistic.

Here we used NST to study visual preferences in the Rainbow Darter (*Etheostoma caeruleum*), a colourful freshwater fish species native to eastern North America. The Rainbow Darter inhabits creeks and small to medium rivers with gravel substrate (Kuehne & Barbour, 2014; Page & Burr, 2011). As for most species in this genus, Rainbow Darters are characterized by visually complex nuptial color patterns, which are expressed only by males, suggesting that these patterns are evolving due to sexual selection. Analyzing the body patterns of ten species in the same genus, including the Rainbow Darter, Hulse and colleagues (2020) found a correlation between the Fourier slope of male body patterns and that of their respective habitats. Female patterns did not show this correlation, suggesting that only the nuptial ornamentation mimics the spatial statistics of the species’ habitat.

We, therefore, aimed to test whether the correlation between the spatial statistics of male luminance patterns and their preferred habitats could be driven by sexual selection. The processing bias hypothesis stipulates that patterns that mimic the spatial statistics of natural scenes are easier to process and thus preferred by conspecifics (Renoult & Mendelson, 2019). Using the style transfer technique, we manipulated the luminance patterns of video animations of male Rainbow Darters to be either more or less similar to their typical gravel habitat (“native habitat” hereafter). This method allowed us to investigate the pattern of a signal holistically, that is, without arbitrarily focusing on an isolated characteristic (e.g. the Fourier slope) as is more traditionally done. We then tested the preference of male and female darters for images of conspecifics that do and do not exhibit the “style” of their native habitat. We predicted that animations of males whose body patterns are the most similar to their habitat’s statistics, thus being easier to process, will be preferred over animations that are less similar to their habitats.

## Methods

### Fish collection and maintenance

We collected male and female Rainbow Darters from Israel Creek (39°29’35.1492’ N, 77°19’28.0488’’ W) and Tuscarora Creek (39°15’6.948’’ N, 77°28’48.288’’ W) in Frederick County, Maryland, between March 15th and April 21st 2022. Fish were transported to the University of Maryland, Baltimore County in aerated coolers and individually housed in a recirculating aquarium system (Aquaneering, Inc.). Tanks contained no substrate to avoid confounding effects but were enriched with a small artificial tree and a bag of crushed coral. Fish were provided natural lighting matching their photoperiod with a constant temperature of 19°C and fed a diet of frozen bloodworms every other day. Finally, males were visually isolated from each other to avoid aggressive behaviours and from females to prevent courtship behaviours but females could see each other. Permission to collect fish was granted by the Maryland Department of Natural Resources Fishing and Boating Services Scientific Collection (permit number SCP202246A). Housing and experimental procedures were approved by the Institutional Animal Care and Use Committee of the University of Maryland, Baltimore County (protocol 623).

### Stimuli creation

We used the Neural Style Transfer (NST), a CNN-based algorithm developed by Gatys and colleagues (2017), to alter the spatial statistics of photographs of male Rainbow Darters. Photographs were taken of males from Middle Fork Red River, Trammel Fork, and Salt Fork in Powell County, KY, Allen County, KY, and Vermilion County, IL, respectively, according to methods in Hulse et al., (2020), using a Canon EOS 5D Mark IV digital camera under standard lighting conditions and Zerene Stacker to combine each stack into an image. Habitat videos were taken using an Ikelite 200DL underwater housing for a Canon EOS 5D Mark IV digital camera equipped with a Sigma 24 mm f/1.4 lens (see Hulse et al., 2020 for details). Rainbow Darters are found almost exclusively in stream habitats with abundant gravel or boulder, and very rarely in open sandy habitats (Page et al. 2011; Kuehne and Barbour 2014). Thus, we considered images of gravel-bottomed streams to represent the “native” habitat and sandy-bottomed streams to represent the “foreign” habitat. Notably, sand habitats are also the most different from gravel habitats when comparing the Fourier slope of freshwater habitat types (Hulse et al., 2020).

A VGG-19 pre-trained network (Simonyan & Zisserman, 2015) with average pooling instead of max pooling and the L-BFGS algorithm for optimisation was chosen for the NST architecture. The content information corresponds to the fourth block of feature layers (out of five) of the CNN while the style information is extracted via a Gram matrix (Figure 1). Briefly, the gram matrix G is the inner product of vectorised (flattened) feature maps F extracted for each layer separately, such that G = F^T^F, and corresponds to the degree of correlations between the feature maps, that is, how often those features co-occur in an image. Such features can be orientations, variations in brightness, patterns, or shapes, and they correspond to the texture of an image.

To render one content image in the style of another image, the algorithm computes and minimises the total loss of the generated image, which corresponds to the weighted sum of the content loss and the style loss such that

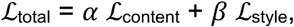

where α and β are the weighting factors for content and style reconstruction, respectively. The content loss is the squared Euclidean distance between the respective intermediate higher-level feature representation F of the generated image (x) and content image (x_c_) at layer ℓ:

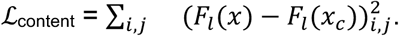

The style function is the sum of the squared Euclidean distance between the gram matrices of the generated image (x) and the style reference image (x_s_) at layer ℓ:

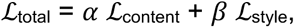

with *w*_ℓ_ as a scaling factor that equals 1 divided by the number of active layers, and *E* calculated as:

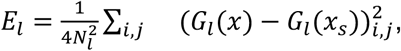

with

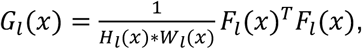

and *H*_l_(*x*) ∗ *W*_l_(*x*) corresponding to the height and width of each feature map at each layer. This last formula corresponds to the simplified version G = F^T^F described above.

For our study purposes, a photograph of an original fish and a photograph of the habitat (either gravel or sand) were fed to the NST algorithm as content and style images, respectively. Content and style have adaptable weights to improve the final result. We kept them constant across stimuli, using the recommended values (content weight = 1; style weight = 1e3). For the style, we only considered the two first blocks of convolutional layers (out of five) as they tend to represent the local statistics of a given image. This means that the scaling factor *w_l_* was equal to 0.5 for layers 1 and 2 and to 0 for layers 3 to 5. For the content, we kept the original parameters and used the feature maps from the convolutional layers of the fourth block.

The original colours of the fish (i.e. the content image) were put back on the final image, using the colour control option of the NST (luminance-only transfer), given that color is known to affect preferences in darter fish (Williams et al., 2013; Williams & Mendelson, 2013). Briefly, the style transfer manipulates the luminance channel only, by first converting both images from the BGR space to the YCbCr space, a space that separates luminance information from colour information. The loss is computed on the luminance channel of that space for both images. The luminance of the output image is then combined with the CbCr information of the input style image, so that the output image keeps the colours of the style image instead of the content image. An example of such output images is shown in Figure 1B.

The PyTorch (Paszke et al., 2019) implementation of the algorithm (https://pytorch.org/tutorials/advanced/neural_style_tutorial.html) with our modifications, as well as the detailed procedure used to create the stimuli, are available on a GitHub repository: https://github.com/yseulthb/NST_fish.

To control for potential biases, we used three images per habitat category (gravel or sand) and styled five different male individuals. Stylized fish were then animated using Blender (Blender Online Community, 2018). The path followed by the animated fish reproduced the path of a live fish, whose video was taken as a representative example of natural motion (Blender files provided on the GitHub project folder). Animations were displayed in underwater videos depicting either gravel (species-specific habitat) or sand (foreign habitat). We selected three different background videos per habitat. For each pair of video animations presented to test individuals, we matched their luminance values (both in terms of background and animated fish). Pairs of videos also presented different animated individuals. Beyond that constraint, all possible combinations of replicates of individual male image, habitat image, and habitat video were randomized across trials and focal individuals.

In summary, we transferred the style of habitats to the body of male Rainbow Darters (one habitat image per fish). Animated images of artificially stylized and nonstylized fish were then presented to live conspecifics, either a male or a female. In addition to manipulating the fish patterns, we also alternated the video background of the fish animations to determine whether individuals prefer the style of fish that is more similar to the background against which it is displayed, regardless of the habitat type (i.e. native vs foreign), i.e., testing a preference for background matching per se. We compared the amount of time fish spent with the animated fish exhibiting the native habitat style (and with a foreign habitat style) when shown in a native habitat vs in a foreign habitat video background. Finally, we explored the influence of style transfer by comparing preferences for stylized vs nonstylized animated fish.

### Dichotomous choice experiments

We ran a series of six experiments to determine whether Rainbow Darters showed a preference for animations stylized with their native habitat statistics. All trials took place during the known breeding season of the species. To avoid habituation, fish were never tested two days in a row, with at least a day between two test sessions. Tested individuals (“focal fish” hereafter) were presented with a pair of stimuli that displayed different animated fish but the same video background. This dichotomous choice experimental paradigm is classically used in the study of mate preferences and has been extensively used in darters to study both male and female mate preferences (Mattson et al., 2020; Roberts et al., 2017; Roberts & Mendelson, 2017). Dichotomous choice tests also show that male darters strongly prefer the nuptial color patterns of other conspecific males, suggesting that male-male competition is also driven by pattern preferences (Williams & Mendelson 2013). The different experimental conditions, their characteristics and sample sizes are summarised in Table 1. In addition to estimating preference for differently styled fish, we estimated in a seventh experiment their preference for their habitat, irrespective of the animated fish (i.e. while showing the same animated, unstyled fish on both sides). This experiment allowed us to test whether individuals have a preference for their native habitats, which is a fundamental assumption of our hypothesis.

**Table 1:** Experimental conditions and sample sizes for the animated fish (Experiments 1 to 6) and the habitat (Experiment 7) preference experiments. Grey cells indicate column names.

| Experiment | Background video | Style of fish 1 | Style of fish 2 | Sample size |
| --- | --- | --- | --- | --- |
| Experiment 1 | native | native | foreign | M: 20 F: 17 |
| Experiment 2 | foreign | native | foreign | M: 17 F: 16 |

| Experiment 3 | native | native | nonstylized | M: 19 F: 13 |
| --- | --- | --- | --- | --- |
| Experiment 4 | native | foreign | nonstylized | M: 16 F: 15 |
| Experiment 5 | foreign | native | nonstylized | M: 20 F: 17 |
| Experiment 6 | foreign | foreign | nonstylized | M: 19 F: 16 |
| Experiment | Fish stimulus | Habitat video 1 | Habitat video 2 | Sample size |
| Experiment 7 | nonstylized | native | foreign | M: 16 F: 13 |

**Table 2:**
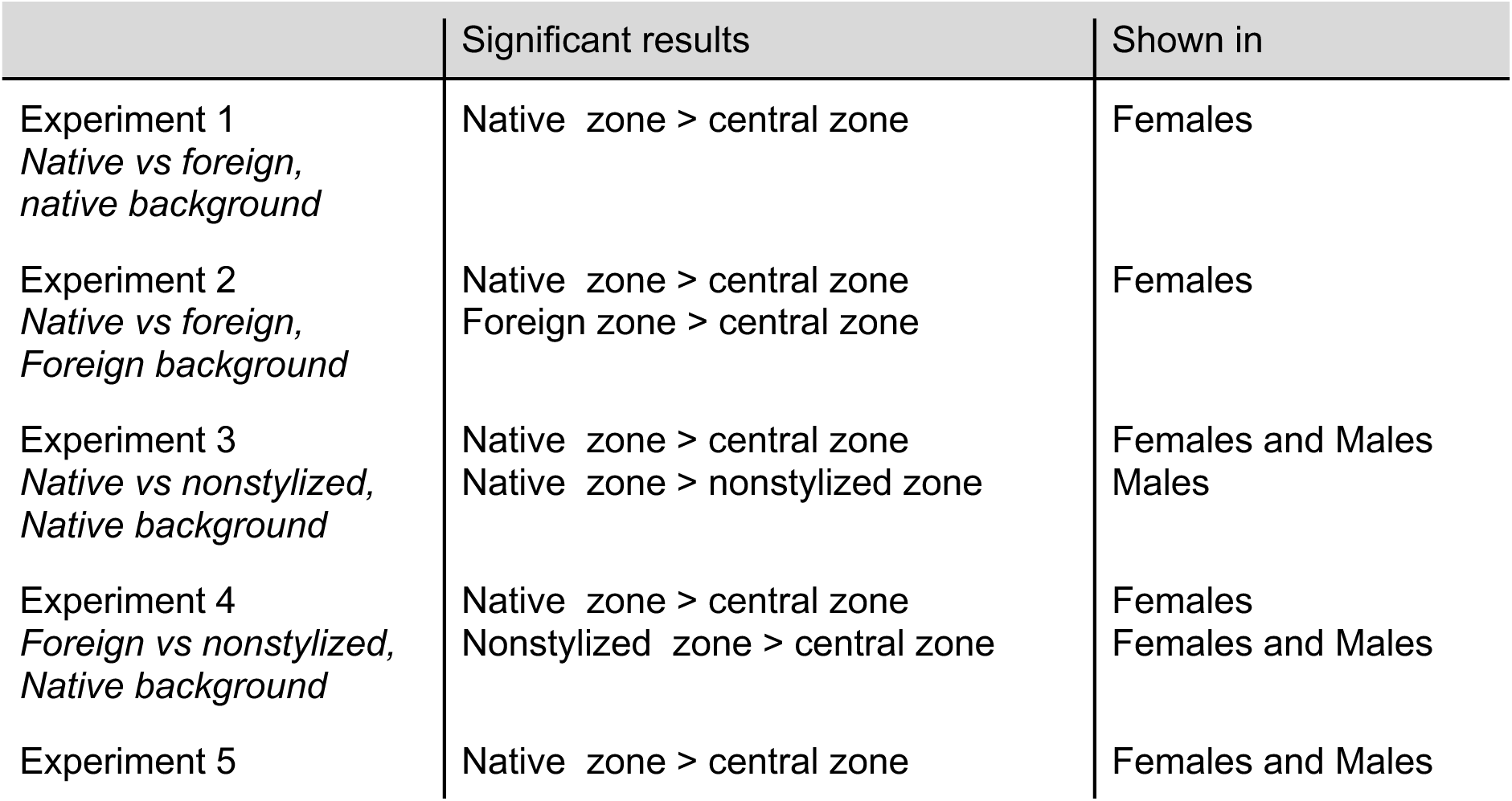
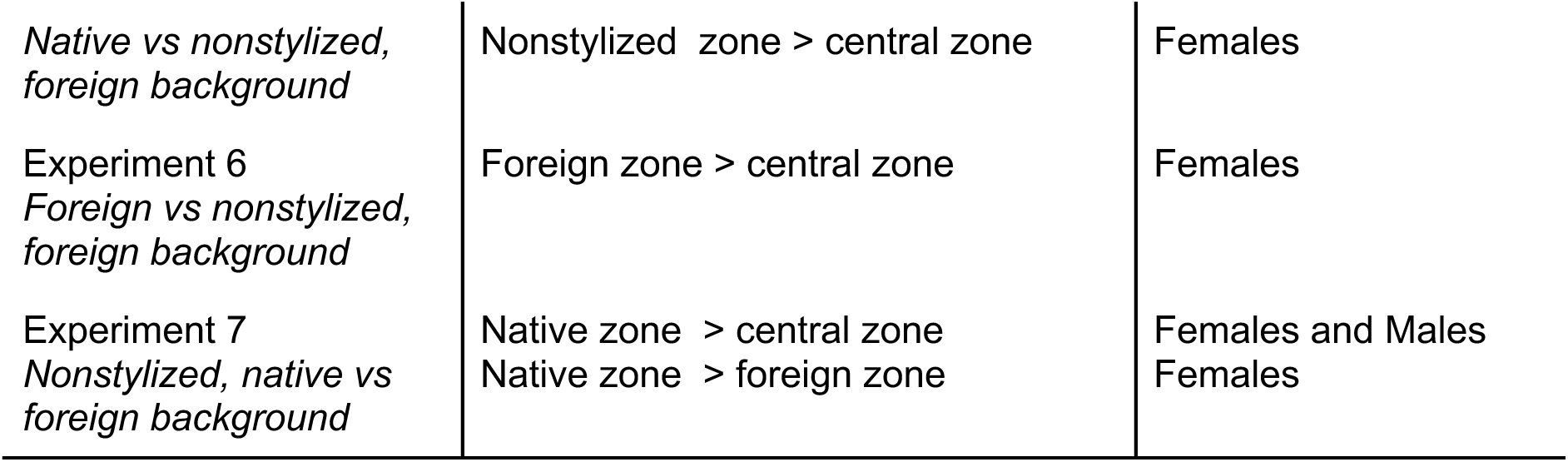
Summary of significant results (2-way ANOVA, *p* < 0.05).

Trials took place in a 37.9L tank flanked by two monitors along the short sides displaying the video animations. The long sides of the tank were covered on the inside with nonreflective black plexiglass to inhibit fish from seeing and interacting with their own reflection. Water was systematically changed between trials and tanks were filled up to 75% of their capacity. Trials were video-recorded using an overhead video camera (Logitech). A 12cm wide zone in the middle of the tank was delineated with transparent removable dividers and served as an acclimation area. Association zones were defined as the 5cm-wide area in front of the video animations (see Figure 2). This distance corresponds to the usual courting distance between males and females (darters are pair spawners) and is in the range of an estimated visual acuity for that species (Caves et al., 2017).

**Figure 2:**
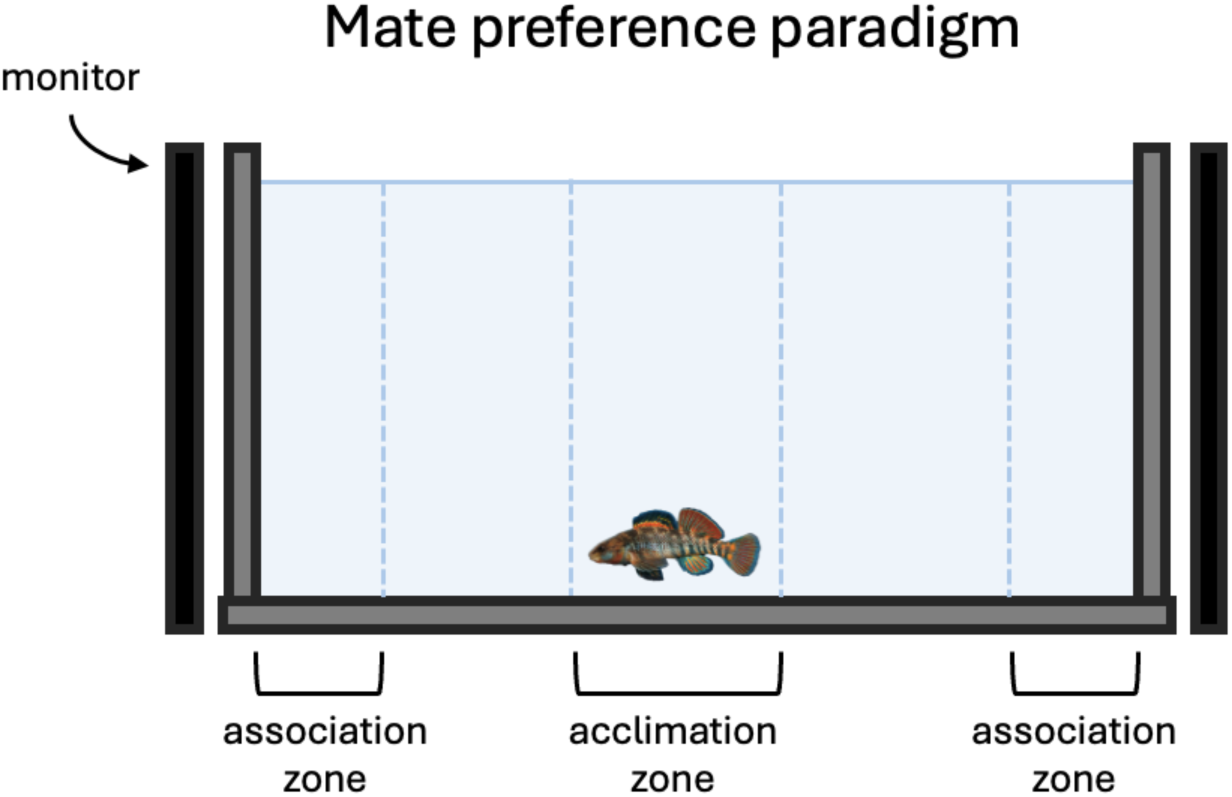
Experimental protocol. The focal fish is shown two different stimuli on monitors situated along the short sides of the tank. The stimulus to which it is more attracted is defined by comparing time spent in the association zones, the 5cm-wide area in front of the video animations.

To control for experimental side bias, each trial alternated on which side of the tank the stimulus conditions (e.g. gravel-styled or sand-styled) were shown. Most fish took part in all seven trials, while the experiment order was randomised between individuals to control for potential order effects. Furthermore, using five individuals as animated fish combined with the different modification styles limits any recognition bias that could influence our results. Additionally, those five animated fish were paired with each other in all possible combinations to avoid a confounding effect of size difference. It should be noted, however, that fish size difference does not necessarily predict the outcome of male fights in that species (*unpublished experiment*).

For the first five minutes of a trial, focal fish were contained in the acclimation area while opaque dividers blocked the video animations. Those opaque dividers were then removed and the fish remained in the acclimation area for an additional five minutes. This protocol gave the focal fish time to visually assess both sides. Finally, the transparent dividers were removed and the focal fish could freely move in the tank for 15 minutes. We identified the preferred stimulus by comparing the amount of time spent in the two association zones during the first 12 minutes after the transparent dividers were removed. Time spent with a stimulus in a dichotomous mate choice trial is a reliable proxy for preference in freshwater fishes (Aspbury & Basolo, 2002; Brooks & Endler, 2001; Lehtonen & Lindström, 2008). Trial videos were scored using BORIS, an open-source event logging software for video and audio files (Friard & Gamba, 2016), by four persons who were blind to experimental conditions. Three mutually exclusive behaviors were defined: the focal fish was either in the central zone (‘center’) or in one of the association zones (‘left’ or ‘right’). An event would start when the fish entered one of the zones and would end when it left that zone to enter another zone. At the end of the video, the durations for all three events (i.e., time spent in each zone) were summed up separately and the data entered into a shared, online table. Once all videos were scored, left and right zones were associated to their respective conditions (e.g., sand or gravel). In order to ensure all four scorers reached consensus on association times, interobserver reliability was first tested on six trials. Times reported for the three zones were compared across the observers and estimated to be 90% similar between scorers (i.e. 90% of reported times did not differ by more than half a second from each other). Trials were then assigned evenly across three authors for scoring.

### Statistical analyses

For all seven experiments separately, we first checked for a potential side bias by comparing times spent in the left vs right association zones irrespective of the stimuli shown. We used either t-tests or Mann-Whitney Wilcoxon tests depending on the data distribution.

To estimate habitat or animation preference, we compared the averaged time spent in the association zone for each condition as well as the averaged time spent in the central zone. This latter measure informs us of overall interest in the stimuli. To compare times spent in those three areas and as data followed a normal distribution (as indicated by normality assessing Shapiro-Wilks tests), we ran one-way ANOVAs. If ANOVA results were significant, we ran posthoc tests (Tukey HSD tests) to identify which pairs of stimuli were significantly different. We also report the number of entries in each association zone, a proxy for mobility. Finally, we counted the number of times the focal fish entered the association zones and used chi-square tests to identify any difference. To determine whether preferences might vary depending on the video background type (native or foreign habitat), we ran t-tests with unequal variance between pairs of experiments where the only difference was the background video type (experiment 1 vs 2, experiment 3 vs 5, and experiment 4 vs 6). All analyses are reported for males and females separately. Statistical analyses were performed with the software R (R Core Team, 2021) using the car package (Fox and Weisberg, 2019).

## Results

### Experiments 1 to 6: Fish preference

We analysed the data of 13 to 17 females and 16 to 20 males (see Table 1) for animation preference. We initially tested a minimum of 15 and a maximum of 20 individuals per experiment. However, individuals that either froze or stayed in the same location without further moving for the entire duration of the experiment were discarded to avoid inflating the time spent in the central area, as these fish provide no information about time spent in the association zones and therefore no information about pattern preferences. We found no side bias for any of the six experiments testing fish preference, neither for females nor males. Detailed test results are provided in the supplementary material.

<u>Experiment 1</u> tested the preference for a fish animation stylized with the statistics of a Rainbow Darter’s native habitat versus a foreign habitat, with a background video of a native habitat (gravel). According to the processing bias hypothesis, fish should prefer the animation with the statistics of their habitat, as their visual system processes those more easily. Results were significant for focal females only (*F*(2,48) = 4.75 *p* = 0.013 for females; *F*(2,57) = 1.89 *p* = 0.16 for males; Fig. 3A). Post-hoc tests found that the amount of time females spent in the association zones of the animation stylized with the native habitat was significantly greater than the amount of time spent in the central zone (*t* = −3.03, *p* = 0.011).

**Figure 3:**
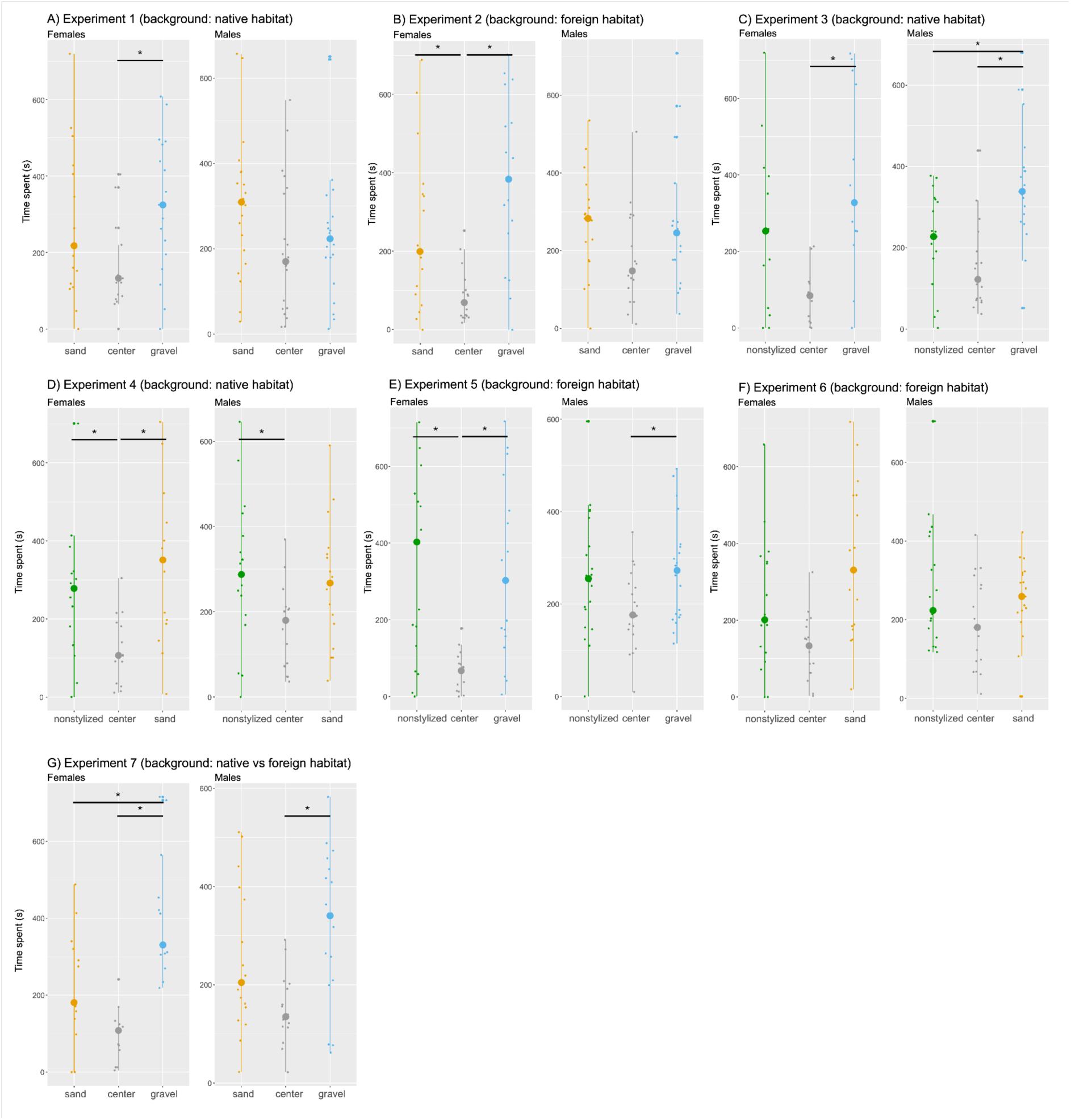
Time (ms) spent in the association or central zones for the seven experiments. Fish animations stylized to reflect the statistics of the species’ native habitat (gravel) are represented in blue, a foreign habitat (sand) in orange, nonstylized animations in green, and the central zone in grey. Asterisks and horizontal bars indicate statistically significant differences between conditions.

On average, males entered the association zone of the fish stylized with the native habitat 6.5 times and of the fish stylized with the foreign habitat 7.8 times. For females, they entered 6.35 times and 5.53 times, respectively. No significant difference was found for either group (X² = 0.1182, df = 1, *p* = 0.731 for males; X² = 0.05708, df = 1, *p* = 0.811 for females).

<u>Experiment 2</u> used the same animation types as Experiment 1 but a background video of a foreign habitat (sand) instead of a native habitat to explore whether the background style influences preferences, as in Menzel et al. (2015). Again, the time difference was significant for females only (*F*(2,45) = 10.3 *p* < 0.001 for females; *F*(2,45) = 10.3 *p* < 0.001 for males; Fig. 3B). Post-hoc tests revealed that females spent more time in the two association zones (native habitat (NH) style: *t* = −4.54, *p* < 0.001; foreign habitat (FH) style: *t* = 2.57, *p* = 0.035), compared to the central zone.

There was no significant difference in the number of entries between the two association zones, for either sex (males: mean NH = 7.1, mean FH = 8.1; X² = 0.07387, df = 1, p = 0.786: females: mean RDH = 4.4, mean FH = 4.2; X² = 0.004106, df = 1, p = 0.949).

<u>Experiment 3</u> compared fish stylized with a native habitat versus a nonstylised (NS) fish, displayed in front of a video of a native habitat. This experiment aimed to determine whether the style transfer procedure influences fish’s preferences. A preference for the styled animation, when it corresponds to the native habitat statistics, should remain present when compared to the natural version of the fish. Results were significant for both males (*F*(2,54) = 13.7 *p* < 0.001) and females (*F*(2,36) = 7.94, *p* = 0.0014; Fig. 3C). Post-hoc tests found that the amount of time females spent in the association zone of the fish stylized with the native habitat was significantly greater than the amount of time spent in the central zone (*t* = −3.96, *p* < 0.001). Males spent significantly more time in the association zone of the fish stylized with the native habitat compared to the central zone (*t* = −5.17, *p* <0.001) and more time in the association zone of the fish stylized with the native habitat compared to the nonstylized fish (*t* = −3.27, *p* = 0.0052). Thus males preferred the fish that was stylized with the native habitat over the natural version of the fish.

There was no significant difference in the number of entries between the two association zones, for either sex (males: mean NH = 6.3, mean NS = 5.2; X² = 0.07387, df = 1, p = 0.786: females: mean NH = 4.9, mean NS = 4.4; X² = 0.004106, df = 1, p = 0.949).

<u>Experiment 4</u> tested a preference for fish stylized with a foreign habitat versus a nonstylized fish, with a background video of a native habitat. A stronger preference for the nonstylized fish (natural phenotype) is expected, as the animation with the foreign habitat statistics should not be better processed by the visual system. Results were significant for both males (*F*(2,45) = 3.45 *p* = 0.04) and females (*F*(2,42) = 6.9, *p* =0.0026; Fig. 3D). Post-hoc tests revealed that females spent significantly more time in the association zone of the fish stylized with the foreign habitat (*t* = 3.65, *p* = 0.0021) and of the nonstylized fish (*t* = 2.44, *p* = 0.0487) compared to the central zone. Males spent significantly more time in the association zones of the nonstylized fish compared to the central zone (*t* = 2.48, *p* = 0.044).

There was no significant difference in the number of entries between the two association zones, for either sex (males: mean FH = 6.3, mean NS = 6.7; X² = 0.07387, df = 1, p = 0.786: females: mean FH = 6.0, mean NS = 6.1; X² = 0.004106, df = 1, p = 0.949).

<u>Experiment 5</u> tested a preference for the animation stylized with the native habitat versus a nonstylized animation, with a background video of a foreign habitat. Similarly to Experiment 3, a preference for the styled animation is expected. Results were significant for both males (*F*(2,57) = 3.52 *p* = 0.036) and females (*F*(2,48) = 10.2, *p* < 0.001; Fig. 3E). Post-hoc tests revealed that females spent significantly more time with the fish stylized with a native habitat (*t* = −3.86, *p* = 0.00099) and with the nonstylized fish (*t* = 3.97, *p* = 0.00073) compared to the central zone. Males spent significantly more time in the association zone of the fish stylized with native habitat compared to the central zone (*t* = −2.43, *p* = 0.048).

There was no significant difference in the number of entries between the two association zones, for either sex (males: mean NH = 7.8, mean NS = 8.0; X² = 0.07387, df = 1, p = 0.786: females: mean NH = 4.1, mean NS = 3.9; X² = 0.004106, df = 1, p = 0.949).

<u>Experiment 6</u> tested a preference for the animation stylized with a foreign habitat versus a nonstylized animation, with a background video of a foreign habitat. Similarly to Experiment 4, a preference for the nonstylized fish (natural phenotype) is expected. Results were significant for females only (*F*(2,45) = 7.24 *p* = 0.0019 for females; *F*(2,54) = 3.01, *p* = 0.058 for males, Fig. 3F). Post-hoc tests revealed that the amount of time females spent in the association zone of the animation stylized with a foreign habitat was significantly greater than the amount of time spent in the central zone (*t* = 3.81, *p* = 0.0013).

There was no significant difference in the number of entries between the two association zones, for either sex (males: mean FH = 8.1, mean NS = 8.7; X² = 0.07387, df = 1, p = 0.786: females: mean FH = 6.6, mean NS = 5.9; X² = 0.004106, df = 1, p = 0.949).

### Background comparisons

A potential influence of the background video on preferences was investigated by showing animations in front of two different backgrounds: one that corresponded to the native habitat of the focal fish and one that showed a foreign habitat. There was no significant difference in time spent in either association zone between experiments that differ only in the type of video background (native or foreign habitat), that is, between experiment 1 and experiment 2 (native habitat style: *t* = −0.481, df = 33.505, *p* = 0.6337; foreign habitat style: *t* = 0.662, df = 34.942, *p* = 0.512), between experiment 3 and experiment 5 (native habitat style: *t* = 0.66602, df = 25.447, *p* = 0.5114; nonstylized: *t* = −0.854, df = 26.827, *p* = 0.4006) and between experiment 4 and experiment 6 (foreign habitat style: *t* = 0.242, df = 24.599, *p* = 0.8105; nonstylized: *t* = 0.194, df = 29.886, *p* = 0.8475). This suggests that background videos did not influence the preferences of focal individuals.

### Experiment 7. Habitat preference

A processing bias hypothesis assumes that animals prefer the visual statistics of their own habitat. To estimate preference for native habitat (NH) versus foreign habitat (FH), we analysed the data of 13 females (out of 15 tested) and 16 males (out of 20 tested). Discarded individuals either froze or stayed in the same location without further moving for the entire duration of the experiment. T-tests for the remaining males and females showed no difference between times spent on either side of the tank (females: *t* = −1.0095, df = 23.992, *p* = 0.3228; males: *t* = 0.79333, df = 29.307, *p* = 0.434); thus we did not exclude any individuals due to side bias.

Results were significant for both males (*F*(2,45) = 6.35 *p* = 0.0037) and females (*F*(2,36) = 17.3, *p* < 0.001; Fig 3G). Tukey’s HSD Test for multiple comparisons found that the amount of time males spent in the association zone of their native habitat was significantly greater than the amount of time spent in the central zone (*t* = −3.54, *p* = 0.0026). Females spent significantly more time associating with their habitat compared to the central zone (*t* = −5.85, *p* <0.001) and more time associating with their habitat compared to the less preferred habitat (*t* = −3.46, *p* = 0.0039). Thus females preferred their “native” habitat over the “foreign” habitat.

There was no significant difference in the number of entries between the two association zones, for either sex (males: mean NH = 6.8, mean FH = 6.5; X² = 0.07387, df = 1, p = 0.786: females: mean NH = 5.4, mean FH = 4.0; X² = 0.004106, df = 1, p = 0.949).

## Discussion

Using the neural style transfer algorithm, we modified animations of male Rainbow Darter fish to reflect the style of (1) their preferred habitat (gravel) and (2) a less preferred (“foreign”) habitat (sand). We then investigated female and male preferences for stylized animations as well as nonstylized animation. We found that, across all experiments, females spent more time in the association zones than in the central zone, but they showed no significant preference for any one pattern over another (native habitat style, foreign habitat style, and nonstylized). In contrast, males spent more time in the association zones in only half of the experiments (Experiments 3, 4, 5); however, when they showed a significant preference, males preferred animated fish stylized to reflect their preferred habitat over the nonstylized animated fish, spending more time in the corresponding association zone than in the two other zones (Experiment 3). As for habitat preferences alone (Experiment 7), females preferred the video of their habitat, spending more time in the corresponding association zone than in the two other zones. Males, on the other hand, only preferred their habitat over the central zone; no difference was found between times spent associating with videos of gravel versus sand habitat.

Our finding that males showed a preference for the fish stylized with the native habitat over a nonstylized fish (Experiment 3) provides some support for sensory drive and the processing bias hypothesis (Endler, 1992; Endler & Basolo, 1998; Seehausen et al., 2008; Renoult & Mendelson, 2019). Sensory drive predicts that animals will have habitat-specific preferences, and processing bias suggests that those preferences are determined by redundant patterns in the environment. As such, processing bias is an example of a pre-existing preference that derives from adaptations of the receiver’s sensory and cognitive systems in other contexts, which can be exploited by signalers to increase mating success (Ryan & Cummings, 2013; Fuller & Endler, 2018). For example, pioneering studies on the Túngara frog revealed that females’ preferences for chucks, a specific component of males’ calls, are influenced by the tuning properties of their auditory system (see e.g. Ryan et al., 1990). In that case, sexual selection favored the evolution of male traits, here the chuck calls, that exploit pre-existing female biases emerging from the property of their auditory system, shaped by their environment (Ryan & Keddy-Hector, 1992). The majority of examples of pre-existing preferences that have been explained in the context of sensory drive have been limited mostly to simple features. Thus, pre-existing preferences for complex patterns, such as the preference for artificial crests in Australian finches (Burley & Symanski, 1998), are often assumed to emerge incidentally or are not well explained. The processing bias hypothesis provides an explanation for pre-existing preferences for complex patterns, and the attraction of males in our study towards fish stylized with the statistics of their native habitat supports that hypothesis. However, it should be noted that this preference does not appear consistently across experiments (e.g., no such effect in experiment 5), nor was it present in females. Nonetheless, other results are at least consistent with expectations of processing bias, with fish spending significantly more time with the native habitat style as compared to the neutral zone. One solution may be to increase the sample size. Studies in humans that demonstrate a pre-existing preference for the spatial statistics of natural scenes typically have sample sizes of up to 50 participants (see e.g. Menzel et al., 2015; Spehar et al., 2015). Reaching such a sample size with wild animals is a challenge, but not an insurmountable one.

Across our experiments, we tested the preferences of both males and females, as both sexes can play a role in sexual selection, and previous research suggests that male and female preferences for a given trait can be either similar or opposed (Candolin, 2004; Qvarnstrom & Forsgren, 1998; Wong & Candolin, 2005). In darters, males compete physically and vigorously with other males for access to females, defending their territory, challenging courting males, and mate guarding, and both sexes are choosy when it comes to selecting a mate. Females and males prefer the same conspecific male color and pattern in at least two species of darters, *Etheostoma barrenense* and *E. zonale* (Williams & Mendelson, 2013), but whether they share a similar preference for the style of the pattern might depend on its function. For example, a male pattern might signal dominance to other males and predict the likelihood of winning a fight with that individual (see e.g. Guerrera et al., 2022); a male might prefer to interact longer with a male against which it would win. Therefore, it would be interesting to compare the pattern of the observing fish to the displayed fish to determine whether certain visual characteristics could predict the preferences of focal individuals. For females, the same pattern could indicate a male’s fitness or paternal quality; females would presumably spend more time associating with a higher-quality male. Only males in our study showed a significant preference for animations with the style of their preferred habitat over a foreign habitat; however, our results do not allow us to determine whether male and female preferences are significantly aligned or opposed. Whether intra- or intersexual processes play a more important role in the design of darter sexual signals therefore remains an open and interesting question.

In addition to manipulating the fish patterns, we also alternated the video background of the animations to determine whether individuals prefer the style of fish that is more similar to the background against which it is displayed, regardless of habitat type (i.e. native vs foreign). This design allowed us to test for a preference for background matching per se. Comparing experiments that differed only in background video (Experiment 1 vs Experiment 2, Experiment 3 vs Experiment 5, and Experiment 4 vs Experiment 6; Figure 4) reveals no effect of background on association time. For each relevant comparison, there were no significant differences in the amount of time spent in each of the three zones, indicating no effect of the background, as that is the only difference between each pair of experiments. This result was somewhat surprising, as a previous study in humans that manipulated the Fourier slope of the background against which faces were presented found that the background slope influenced the attractiveness of the faces (Menzel et al 2015). Sample sizes were higher in the human study than in ours, but an additional explanation is that the Fourier slope has a different effect on preferences than does the style of an image.

### Limitations and future directions

The camera viewpoint is an important factor to consider when choosing style images. We chose wide-angle images from the perspective of a benthic fish, keeping the camera close to the stream floor. Wide-angle images that include an entire visual scene are usually the go-to when testing the adaptation of the visual system to natural scene statistics (e.g. Gupta et al., 2023). Indeed, the efficient coding hypothesis states that the visual system is adapted to such statistics (Olshausen & Field, 1996; Simoncelli & Olshausen, 2001), but it does not necessarily specify other aspects of visual perception, such as visual acuity or the breadth of the visual field of a given species. However, at least one study, of mice, was able to predict visual perception with cameras filming the environment from the perspective of those animals (Qiu et al., 2021). We assume that similar retinal adaptation takes place in darters and thus we used a similar approach, using images taken from the perspective of the animal.

The scale of the style images might also affect relevant statistics. For instance, habitat images are not scale invariant; thus, the scale of the image affects which image properties are transferred (e.g., more or less fine-grained details, more or less repetitive patterns) and thus the preference of the observer fish. Experimenting with different scales would enable direct testing of how scale changes the properties of the output image and the preferences of the fish. In addition, investigating the image properties of different stylized images could help disentangle which properties influence fish preferences, from their Fourier slope (as mentioned on Figure S3) to their contrast value. In the current study, only the mean luminance (i.e. brightness) difference was minimized between pairs of stimuli; thus, it remains possible that the observer fish noticed differences in an unknown image property that influenced its visual preference. Follow-up studies should consider aspects such as image scaling and visual acuity.

In our experiments, we kept the color of the animated fish constant. One important future direction will be to test the interaction between colors and patterns while modulating spatial statistics of signal designs and ask how this influences preferences. Another future research direction will be to focus on specific body parts of the fish rather than the body as a whole, to assess whether some parts are more susceptible to sexual selection. For instance, Gumm & Mendelson (2011) found no correlation between body colors and fin colors of several darter species, suggesting that patterns on these different body parts may be evolving independently. These differences make sense since dorsal fins are often erected for courtship or male-male fights (see e.g. Johnston, 1994) and concealed at rest. Fin patterns therefore may be evolving due to sexual selection alone and be less constrained by camouflage. Conveniently, the most recent implementation of the NST algorithm (Gatys et al., 2017) has two control options that make it useful for addressing such questions: spatial and color controls. The spatial control lets the user predefine on which part of an image to transfer the style. This is useful for modifying subparts of a body and identifying potential differences in their effect on visual preferences or communication. With color control, the model allows the user to either retain the original color of the content image (i.e., to “transfer” only achromatic information) or to transfer both chromatic and achromatic information. This option is convenient for testing the influence of color on visual preferences.

Other methods could also be used instead of the current implementation. For instance, the Laplacian Pyramid method (Burt & Adelson, 1983; Sunkavalli et al., 2010) enables the transfer of texture information (the “style” of an image) from one image to another and represents a less computationally intensive alternative. We also recommend exploring alternative and ever-evolving Neural Style Transfer methods. For instance, the comparative work of Wright & Ommer (2022) could guide the reader in determining which method would lead to better results both in terms of qualitative output and required computational power.

Our implementation, based on a PyTorch tutorial, makes it convenient to style any new image even for an audience not familiar with the language or the concept of neural style transfer. The code, in the form of a notebook, only requires a few adaptations, such as changing the weights put on the style and the content image depending on the desired output. Comments in the code should help to run the script easily even without prior knowledge of deep learning algorithms. Finally, because the algorithm is based on the VGG-19 architecture pretrained on ImageNet, a generalist database containing images of diverse objects, people and animals, no additional training is required for the algorithm, making it more straightforward to implement for any new pair of images.

Our study and suggestions for future research demonstrate how the style transfer algorithm and other AI-based techniques can be used to study the evolution of signal design. For instance, the ability of style transfer via the Gram matrix to separate texture information makes it particularly suitable to study camouflage, as highlighted in Hulse et al. (2022). Specifically, using style transfer, one can distinguish between two camouflage strategies: background matching, which depends on texture (i.e., patterns and variations in color or shading) to blend in with the environment, and disruptive coloration, which hinders individual recognition by breaking up a visual outline, thereby creating visual confusion. Using a similar approach in fish, Hulse et al. (2022) compared the gram matrices of images of male and female darters to habitat images of different categories to determine whether fish species were more visually similar to the habitat in which they occur as compared to habitats in which they are typically not found. Female patterns were found more similar to their habitats than were nuptial male patterns, highlighting a potential trade-off between sexual and natural selection (i.e., camouflage) for males during the mating season. Interestingly, that study also found differences between network layers, revealing how those can be used to estimate the influence of spatial scales on the studied trait.

### Conclusion

Using artificial intelligence methods to test the effect of low-level spatial patterning on the visual preferences of fishes, our study provides one of the first attempts to use rapidly advancing AI techniques to directly query animal behavior. Our results bring no clear support to the prediction that male nuptial patterns matching the visual statistics of their habitat are preferred over patterns matching a foreign habitat or a nonstylized fish. However, the hypothesis of a processing bias is also not rejected. Indeed, the significant results we obtained are either consistent with or support the hypothesis. In particular males significantly preferred animations whose patterns match the native habitat over nonstylized animations. Importantly, our study demonstrates how artificial neural networks can be used to test hypotheses about animal signal evolution, by creating or manipulating stimuli without the need to target a specific feature. Indeed, such generative models will allow us to move toward a more holistic perceptual representation of complex signals, paving the way to exciting new questions.

## Data Availability

The PyTorch implementation of the algorithm with our modifications, as well as the detailed procedure used to create the stimuli, are available on a GitHub repository: https://github.com/yseulthb/NST_fish, as recommended by Huettmann & Arhonditsis (2023).

## Acknowledgements

We would like to thank the students of the DART lab who helped with trials and video scoring. This work was supported by the National Science Foundation grant NSF IOS 2026334.

## Supplementary material

**Figure S1:**
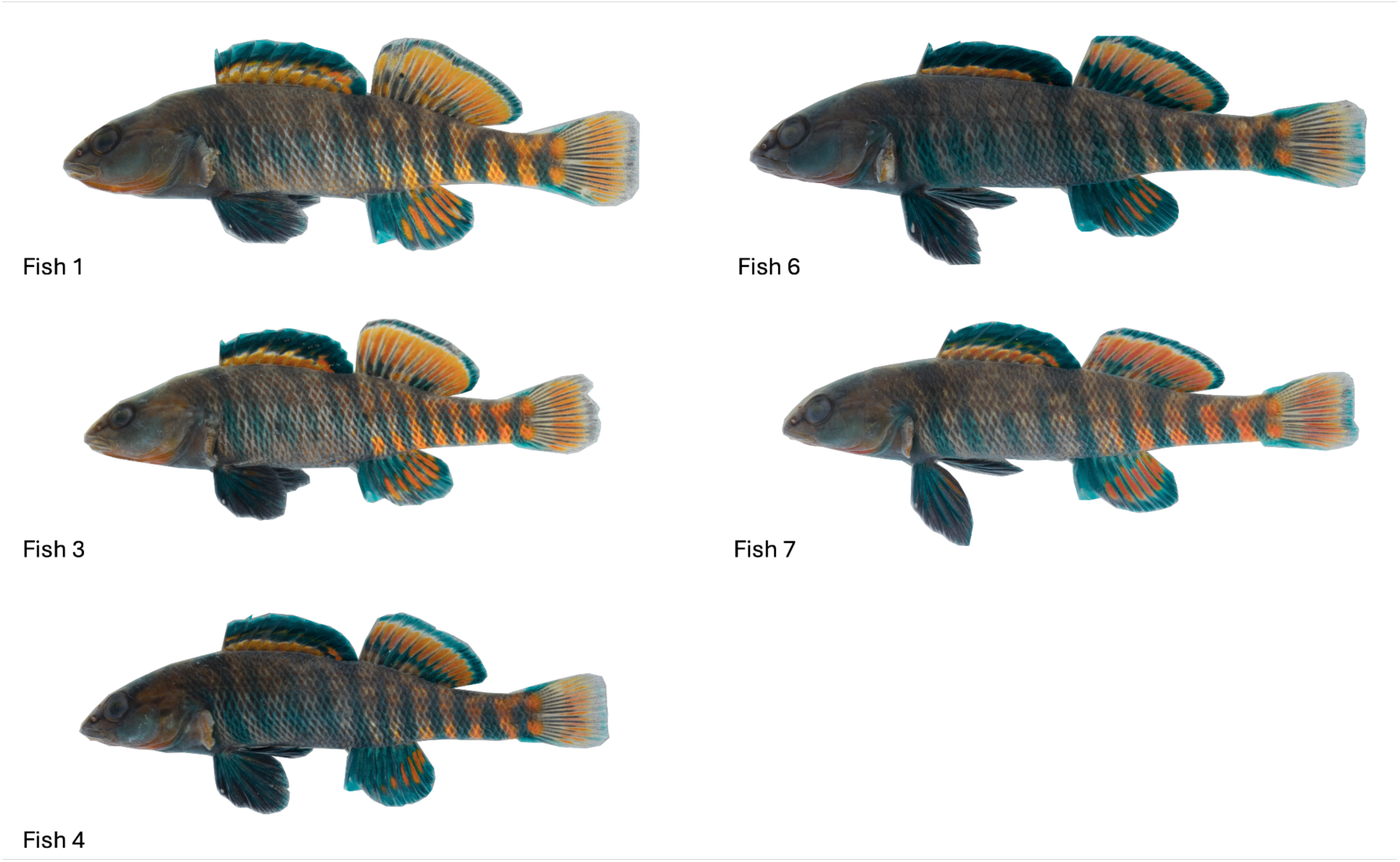
Fish images used as content images.

**Figure S2:**
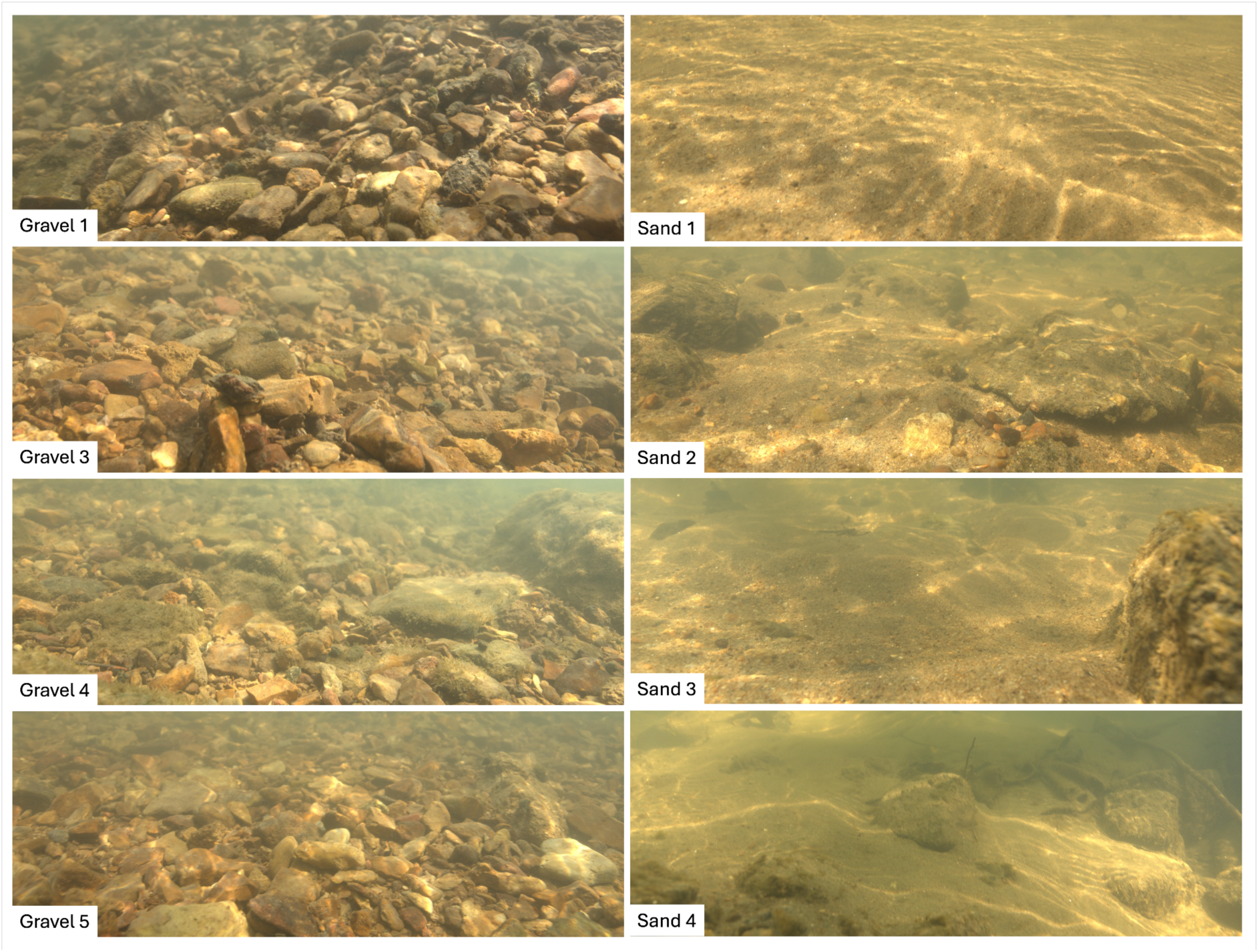
Underwater images used as style images.

**Figure S3:**
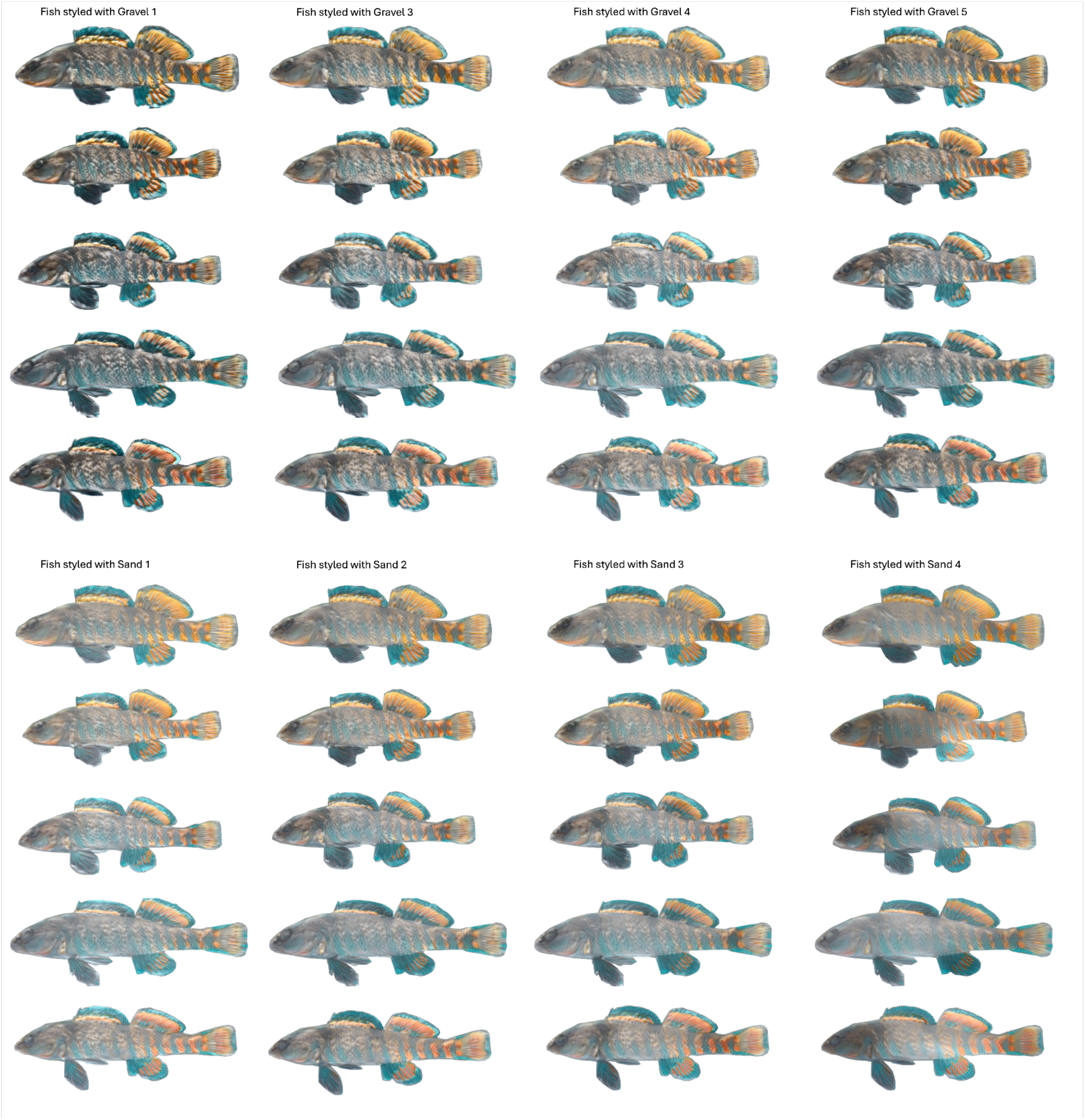
Stylized fish.

**Supplementary table.** Assessing potential side bias in the seven experiments.

|  | Males | Females |
| --- | --- | --- |
| Experiment 1 | $t = 1.765$, $df = 38$, $p = 0.0855$ | $t = -1.758$, $df = 31.81$, $p = 0.0883$ |
| Experiment 2 | $t = 1.952$, $df = 26.82$, $p = 0.0614$ | $W = 94$, $p = 0.207$ |
| Experiment 3 | $t = 0.9881$, $df = 35.97$, $p = 0.33$ | $t = 0.07073$, $df = 23.98$, $p = 0.944$ |
| Experiment 4 | $t = -0.3773$, $df = 26.28$, $p = 0.709$ | $t = -1.626$, $df = 27.84$, $p = 0.115$ |
| Experiment 5 | $t = 1.071$, $df = 37.92$, $p = 0.291$ | $W = 135$, $p = 0.76$ |
| Experiment 6 | $t = 0.2342$, $df = 34.42$, $p = 0.816$ | $t = 0.834$, $df = 29.92$, $p = 0.411$ |
| Experiment 7 | $t = 0.7933$, $df = 29.31$, $p = 0.434$ | $t = -1.01$, $df = 23.99$, $p = 0.323$ |

## Text S1: Fourier slope of stylized images

Although pairs of stylized fish that were presented to fish observers were luminance-matched, they differed in their Fourier slopes. Fourier slopes of fish stylized with the gravel habitat pictures (−2.74 ±0.08) were significantly different from Fourier slopes of fish stylized with the sand habitat pictures (−3.00 ±0.10; t-test: *t* = −8.819, *p* < 0.0001).

